# Amygdala circuit mechanisms underlying alcohol seeking

**DOI:** 10.1101/2024.06.03.596970

**Authors:** Junghyup Suh, Amanda L. Pasqualini, Maria A. Zambrano, Kerry J. Ressler

## Abstract

Alcohol seeking during abstinence is mediated in part by strong associations between the pharmacological effects of alcohol and the environment within which alcohol is administered. The amygdala, particularly the basolateral amygdala (BLA), is a key neural substrate of environmental cue and reward associations since it is involved in associative learning and memory recall. However, we still lack a clear understanding of how the activity of molecularly distinct BLA neurons is affected by alcohol and encodes information that drives environmental cue-dependent, alcohol-related behaviors. We previously demonstrated that a subset of BLA neurons which express the CaMKII and Thy1 markers project preferentially to the nucleus accumbens (NAcc), rather than the central amygdala; and these neurons mediate fear inhibition rather than fear acquisition or expression, suggesting a specific role in positive valence processing. We now demonstrate that Pavlovian conditioning with alcohol administration increases the activity of these Thy1-expressing (Thy1+) excitatory neurons in mouse BLA, which is necessary for the conditioned appetitive response. *In vivo* calcium imaging indicates that the temporal activity profile of these neurons is also correlated with alcohol seeking behavior in response to environmental cues. Optogenetic inhibition of BLA Thy1+ neuronal activity disrupts both the formation and recall of alcohol conditioned place preference. Furthermore, selective axonal inhibition of BLA-Thy1+ neurons reveals that the activity of their NAcc and prefrontal cortex (PFC) projections are differentially necessary for alcohol cue *association* vs. *recall*, respectively. Together, these findings provide insights into a molecularly distinct subset of BLA neurons that regulates environmental cue-reward associations and drives alcohol seeking behaviors in a projection-specific manner.

**Disclosures:** KJR has received consulting income from Acer, Bionomics, and Jazz Pharma; serves on Scientific Advisory Boards for Sage, Boehringer Ingelheim, Senseye, the Brain and Behavior Research Foundation, and the Brain Research Foundation, and he has received sponsored research support from Alto Neuroscience. None of this work is directly related to the work presented here.

## Introduction

During repeated drug use, the processes that link a neutral stimulus, such as environmental cues, with the rewarding effects of the drug can have robust and long-lasting influences on subsequent drug-seeking behaviors ^1, 2^. Alcohol use disorder (AUD) remains one of the most prevalent and important contributors to morbidity and mortality ^3, 4^; yet therapeutic tools for prevention or treatment of AUD are quite limited, and improved understanding of the neurobiology of alcohol seeking behavior is much needed.

Like other potential drugs of abuse, it has been show that when the administration of alcohol is repeatedly paired with different cues, re-exposure to those cues can reinstate alcohol seeking as a result of Pavlovian drug-cue learning ^5^. In clinical studies, motivational reactivity to alcohol-related visual cues has been demonstrated and is thought to be a key mechanism involved in facilitating alcohol craving and eliciting relapse after long periods of abstinence or extinction ^6, 7^. While mesolimbic dopamine broadcasts a general reward signal that likely support the association processes, converging evidence also suggests that multiple memory systems, including the amygdala, NAcc and PFC, contribute to reward-based associative learning and integrate the necessary sensory information in parallel ^2, 8^.

The amygdala is crucial for the acquisition and expression of a variety of motivational tasks, both aversive and appetitive, by providing a neural representation of environmental sensory cues associated with previous drug use and conveying it to downstream brain regions ^9,10^. Human neuroimaging studies demonstrate that exposure to alcohol cues elicits amygdala activity correlated with the motivation to consume alcohol ^11^. Similarly, preclinical studies also indicated that the amygdala is a primary brain region crucial for development of alcohol seeking behavior. For example, environmental cues paired with systemic ethanol injection in a Pavlovian conditioning paradigm increased c-Fos protein expression, a proxy of neuronal activation, in the amygdala ^12^. In addition, olfactory discriminative stimuli, signaling availability of ethanol in an operant self-administration task, also led to a trend towards increased numbers of c-Fos+ cells in the amygdala ^13^. Furthermore, the inactivation of the amygdala, particularly BLA, attenuates Pavlovian-conditioned alcohol seeking behaviors in rats ^14^.

Notably, a previous study with electrophysiological recordings in behaving mice identified at least two populations of BLA neurons associated with differential fear responses ^15^. Specifically, one population (Fear-On) responded to the conditioned stimulus (tone) with increased firing rates during fear conditioning and decreased firing during extinction training, whereas the other population (Fear-Off or Extinction neurons) only responded to the tone with elevated firing frequency during fear extinction. These data suggested at least two distinct functional subpopulations of BLA neurons responsible for fear-related behaviors. Recent studies also demonstrated that subsets of BLA neurons segregated by spatial location in the anterior-posterior axis and downstream connectivity differentially participate in valence-specific behaviors ^16, 17^. However, differentially characterizing behavioral functions of BLA neurons have been hindered by the lack of predictable molecular markers for cell types and their dense differential connections between the BLA and other brain areas, including the NAcc and PFC.

BLA neurons labeled by thymus cell antigen 1 (Thy1) promoter-driven transgenes, such as yellow fluorescent protein (YFP) or Cre recombinase, represent a promising entry point for addressing these issues ^18^. In previous studies, we have extensively examined the BLA Thy1-expressing (Thy1+) neurons using a combination of transgenic mice and viral vectors to express proteins in BLA- and Cre-specific manners ^19, 20^. Our findings suggest that Thy1+ excitatory neurons in the BLA represent a functionally and anatomically consistent subpopulation across transgenic lines; furthermore, they may represent a valence-specific, ‘Fear-Off’ BLA neuronal population that preferentially projects to the NAcc rather than the central amygdala, suppresses fear, and supports fear extinction. However, if these neurons are also ‘Appetitive-On’, representing positive valence cues, and whether they are involved in drug or alcohol-related conditioning behaviors is unclear. More specifically, it is unknown whether exposure to alcohol administration alters the activity of these populations of neurons and how the subpopulations of the BLA contribute to alcohol-cue associations or alcohol seeking behaviors.

We now demonstrate that, in addition to their fear-inhibition properties, BLA Thy1+ neurons are critical for formation and recall of appetitive alcohol-cue associations. The activity of these neurons increased during repeated pairing of alcohol and environmental cues. Using fiber photometry imaging of calcium activity, we show that entering an alcohol-paired compartment coincided with increased activity of Thy1+ neurons, whereas exiting an alcohol-paired compartment concurred with decreased activity. We then used optogenetic manipulation in cell bodies to demonstrate that the activity of BLA Thy1+ neurons is necessary for both formation and recall of alcohol conditioned place preference (CPP). In addition, we found that the activity of their projections to the NAcc and PFC differentially underlies formation vs. recall of alcohol-cue associations, respectively. The findings from this study provide significant support for the notion that the amygdala generates neural representations, via its distinct neuronal populations and their distinct connections, to form distinct positive vs. negative associative memories, and targeting these pathways may provide a novel and robust means of interfering with drug-seeking behaviors.

## Results

### Neuronal activity of BLA Thy1+ neurons responding to repeated pairing of alcohol and context

Since alcohol-mediated CPP depends on learning and memory processes associating alcohol exposure and context information, mice were subjected to a CPP procedure in which B6 mice consistently developed robust CPP responses (Figure 1a). As shown in previous literature ^21, 22^, mice conditioned with alcohol (saline/EtOH or S/E) spent significantly longer time in a compartment associated with alcohol injection (Figure 1b, c), indicating that alcohol had induced a place preference via an association between the context and the rewarding effects of alcohol.

**Figure 1.**
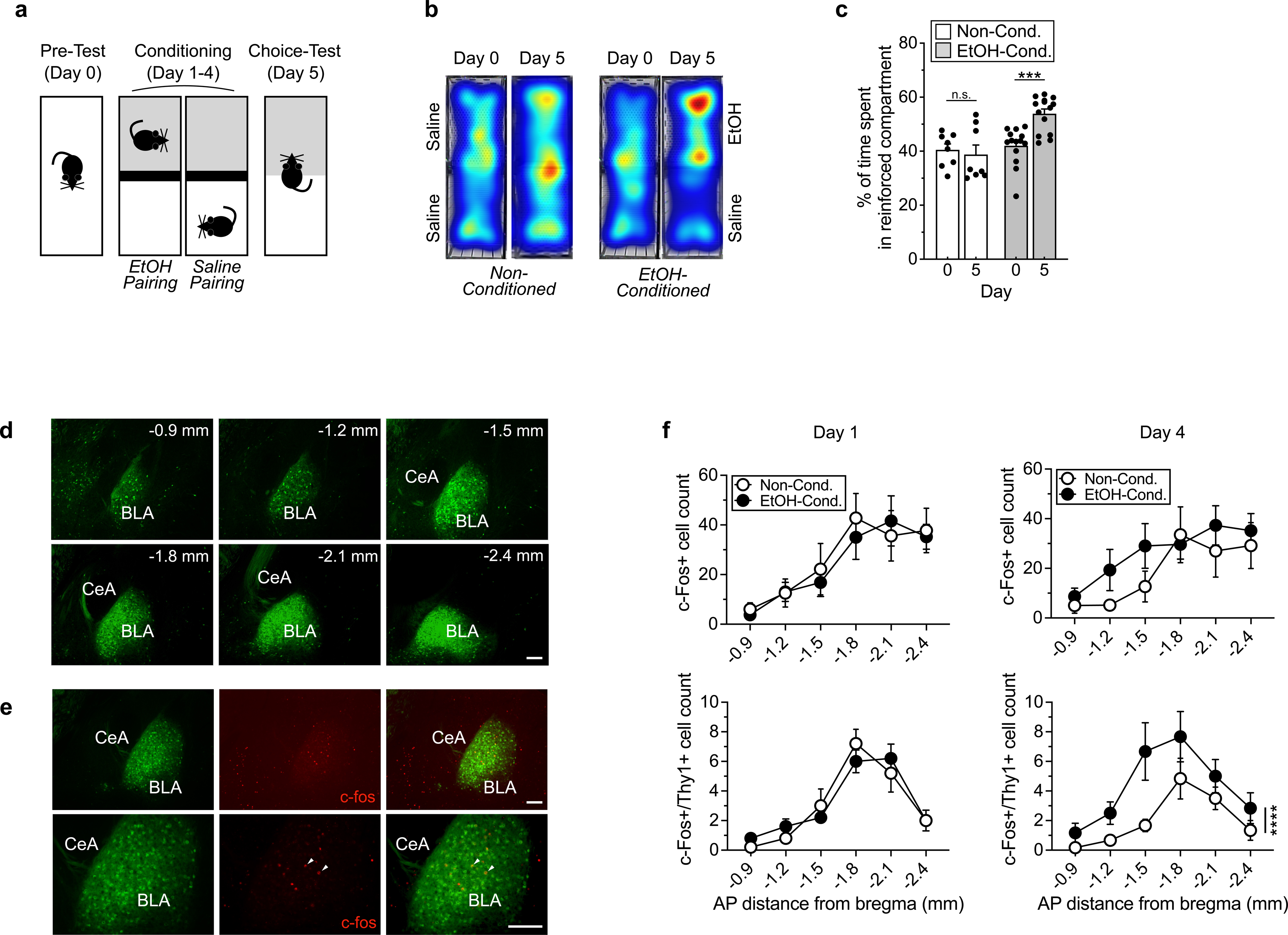
| Ethanol (EtOH)-induced place preference. **a**, Timeline of experimental design. **b,** Heat maps shows time spent in each compartment of the chamber during Pre-Test on Day 0 and Choice-Test on Day 5. **c,** EtOH-conditioned animals formed a preference for the reinforced compartment associated with EtOH administration while un-conditioned, saline-injected in both compartments, animals did not display any preference during the EtOH-free, Choice-Test on Day 5. **d,** Distribution of Thy1+ neurons across AP axis (coronal distance from bregma -0.8 mm to - 2.8 mm) of the BLA. Scale bars: 200 um. **e,** c-Fos expression across the AP axis (coronal distance from bregma of the BLA in response to EtOH administration. **f,** Quantification of c-Fos, eYFP, and c-Fos/eYFP-expressing cells in the BLA.

In contrast, non-conditioned mice (saline/saline or S/S) spent equal time in both compartments (Figure 1b, c). We hypothesized that if Thy1+ neurons represent positive valence-encoding neurons of the BLA, then alcohol administration that elicits alcohol CPP behavior would also activate these neurons. To address this, by capturing dynamic time- and cell type-dependent changes in neuronal activity over the time course of conditioning with alcohol, we subjected Thy1-YFP transgenic mice to the same CPP procedure. The animals were sacrificed 90 min after the end of alcohol conditioning session on Day 1 or 4. All brain tissues were processed to detect c-Fos expression, a proxy for neuronal activity, using immunohistochemistry and the number of c-Fos+ neurons was quantified across the AP axis of the BLA (Figure 1d, e). We found that the total number of c-Fos expressing neurons was greater in the posterior BLA in both non-conditioned and EtOH-conditioned mice on Day 1 and 4. In addition, there was no significant difference in the number of total c-Fos+ cells and that of c-Fos+ cells among Thy1+ cells in the BLA in both groups on Day1. However, the numbers of total c-Fos+ cells and c-Fos+ cells among Thy1+ cells were significantly higher in the EtOH-conditioned group than non-conditioned group on Day 4 of conditioning (Figure 1e). There were approximately 212% more Thy1+ cells also labeled as Fos+ in the ETOH-conditioned group (73 in total from 6 mice) on Day 4 compared to the non-conditioned group (155 in total from 6 mice). The findings indicate that repeated alcohol-context pairings increase BLA Thy1+ neuronal activity, which may mediate positive valence coding.

### BLA Thy1+ neuronal activity is greater when entering an EtOH-paired than a saline-paired context

To examine real-time activity of BLA Thy1+ neurons during associative learning and memory processes for alcohol rewards, we next performed *in vivo* fiber photometry recordings of BLA Thy1+ population activity (Figure 2a). To this end, we expressed a genetically encoded calcium indicator, GCaMP6f, in BLA Thy1+ neurons by injecting a Cre-dependent AAV-Ef1a-DIO-GCaMP6s into the BLA of Thy1-Cre driver transgenic mice and placing an optic fiber above the injection site 3 weeks after virus injection (Figure 2b). We then measured photometry signals during the Pre-Test and Choice-Test to assess the role of BLA Thy1+ neuronal activity in the expression of EtOH CPP behavior that relies on the recall of association between alcohol experience and context (Figure 2c, d). A paired t test revealed a significant increase in preference for the EtOH-paired compartment during Choice-Test compared with the Pre-Test (Figure 2d). Transitions of mouse trajectories between the saline- and EtOH-paired chambers were time-stamped and aligned to photometry signals, and the 8 second window centered around each transition was isolated for analysis (Figure 2e). We observed a transient elevation in BLA Thy1+ neuronal activity starting shortly before and during EtOH-paired compartment entry in EtOH-conditioned mice. A two-way repeated measures ANOVA revealed a significant compartment x test interaction for BLA Thy1+ neuronal activity during entry to the EtOH-paired compartment, with approximately a 99% increase in EtOH-paired cell activity upon entry (Figure 2e, *Inbound*). Conversely, weaker Thy1+ activity during entry to saline-paired chamber was observed. Upon exiting the EtOH compartment, a statistically significant decrease (approximately 79%) in Thy1+ activity was seen (Figure 2e, *Outbound*), upon leaving the conditioned compartment, compared to no relative decrease in controls. These results suggest that the activity of BLA-Thy1+ neurons is spatially correlated with alcohol seeking behavior in response to environmental context.

**Figure 2.**
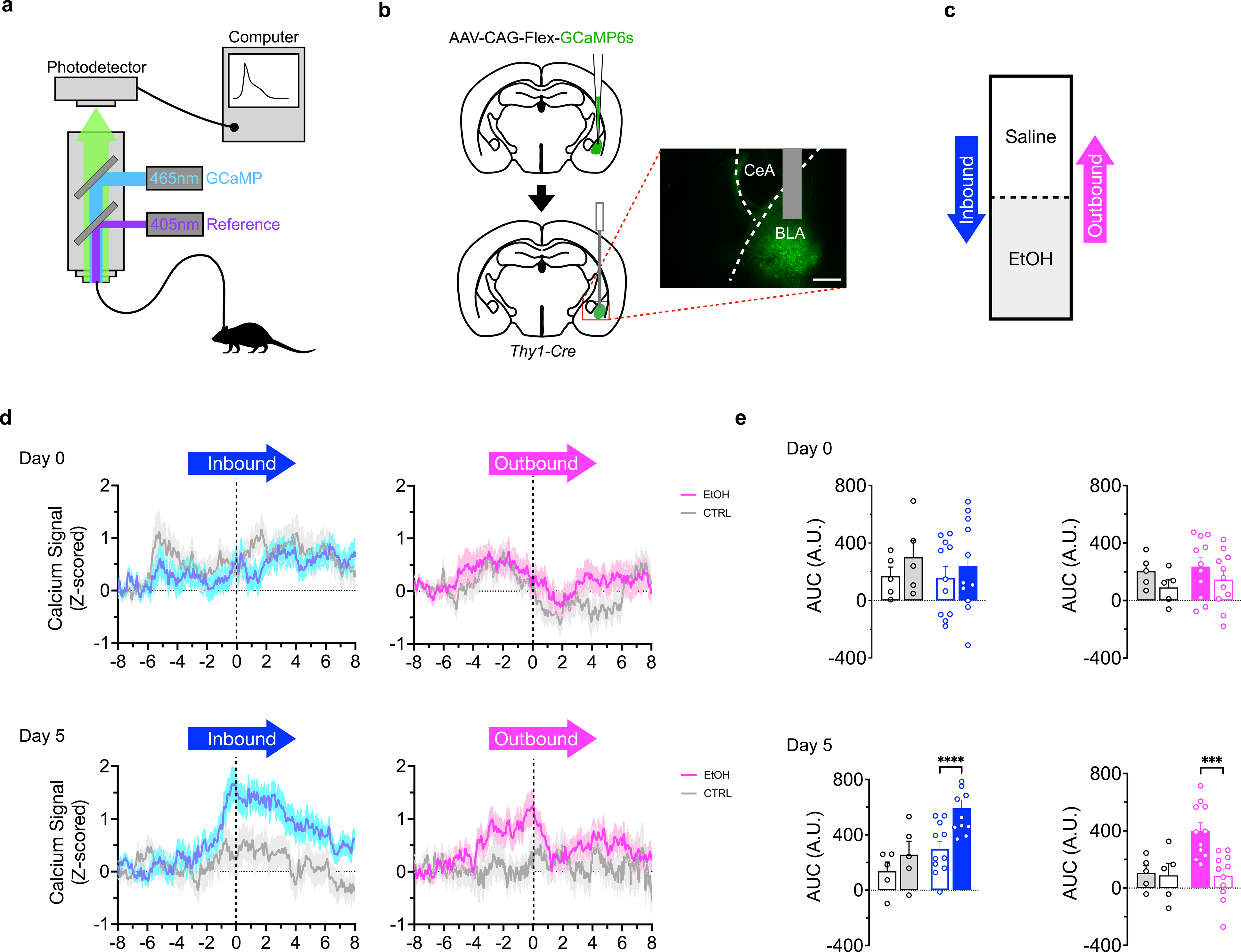
| Fiber photometry of neuronal dynamics during CPP Conditioning and Choice-Test. **a**, Cartoon schematic of fiber photometry setup for recording from mouse. Light path for fluorescence excitation and emission is through a single 400 um fiber optic implantation in the BLA. **b,** Viral expression of AAV-DIO-GCaMP6s and placement of the fiber optic probe in the BLA. **c,** Schematic representation of animal’s trajectories in the CPP box. **d,** The temporal profiles of fluorescent calcium traces during transition between EtOH- and saline-paired compartment entry. Data are illustrated as the z-scored percent change in fluorescence. **e,** Quantification of the fluorescent peaks during transition from one to the other compartment.

### BLA Thy1+ neuronal activity is required for the acquisition and expression of alcohol CPP

To determine if the activity of BLA Thy1+ neurons is necessary for the associative learning and memory processes underlying EtOH CPP, we first tested effects of optogenetically inhibiting BLA Thy1+ neurons. An adeno-associated virus (AAV) directing Cre-dependent expression of the inhibitory opsin halorhodopsin (eNpHR3.0) or control eYFP was bilaterally injected into the BLA of Thy1-Cre mice; both groups of mice were bilaterally implanted with optic fiber ferrules delivering concurrently illuminated 532-nm light during EtOH pairing sessions throughout Conditioning (Figure 3a, b). We found that photoinhibition of BLA Thy+ neurons during conditioning significantly reduced time in the EtOH-administered compartment, indicating a deficit in – or possibly a complete prevention of – the learning and development of place preference (Figure 3c).

**Figure 3.**
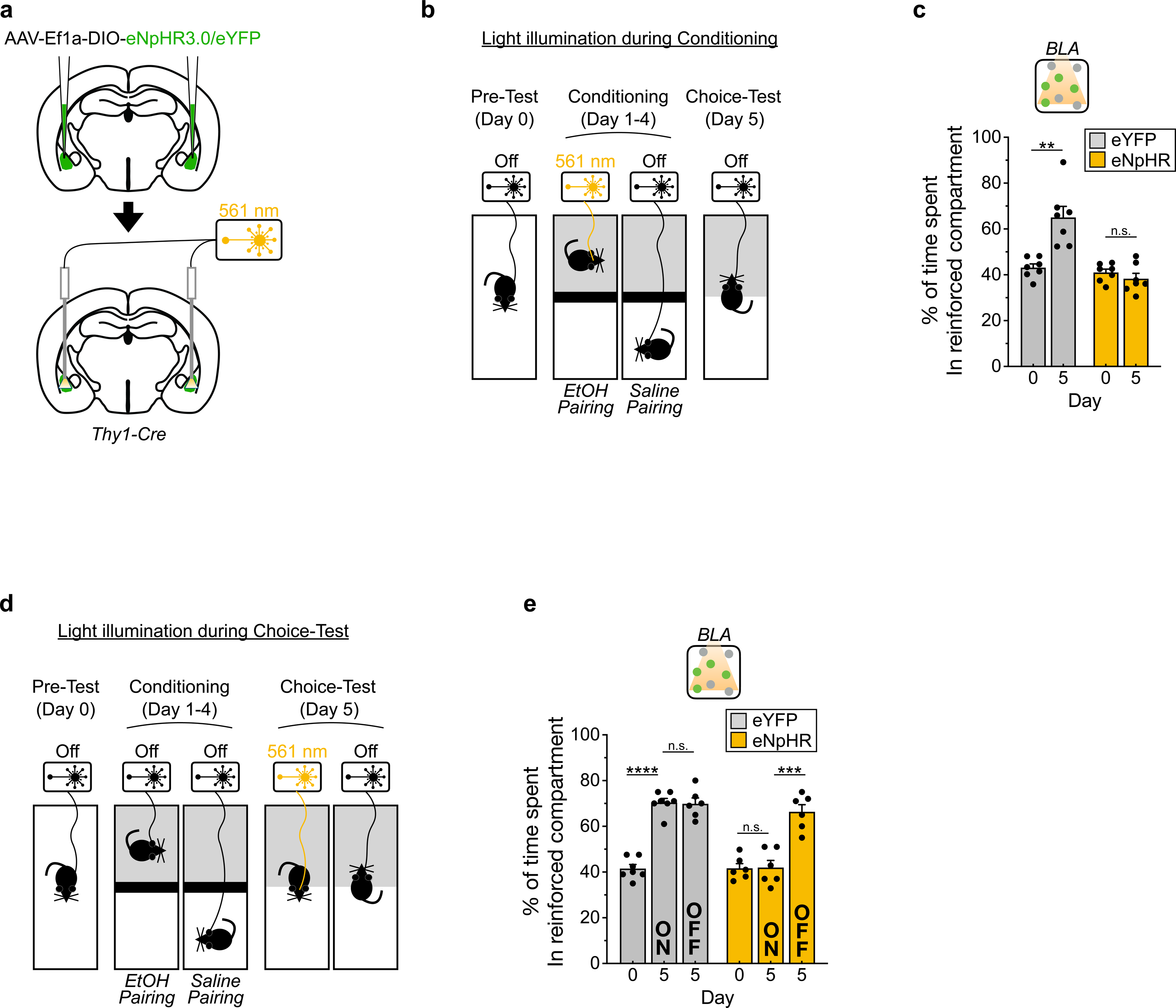
| Thy1+ neuronal activity at cell bodies in the BLA during Conditioning and Choice-Test is necessary for CPP formation and recall, respectively. **a**, Scheme for Cre-dependent eNpHR/eYFP expression in BLA Thy1+ neurons and optogenetic manipulation. **b,** Illumination of light (561 nm, 15-20 mW) started 5 sec before EtOH injection and continued during the entire EtOH-pairing sessions on Conditioning Day 1-4. **c,** Optogenetic inhibition of BLA Thy1+ neurons during Conditioning disrupted CPP formation. **d,** During Choice-Test, light illumination was carried out similarly for the entire session. **e,** Optogenetic inhibition (ON) of BLA Thy1+ neurons during Choice-Test disrupted CPP recall. In a subsequent Choice-Test session without light illumination (OFF), mice injected with eNpHR displayed a similar level of CPP recall compared to mice injected with eYFP.

Since sets of BLA neurons have been shown to express fear memory formed by the association of auditory cues and foot shocks, we separately tested if the activity of these neurons is required for CPP expression or recall after an association of context and alcohol experience has already been formed. We inhibited BLA Thy+ neuronal activity in separate cohorts of mice only during the Choice-Test (Figure 3d). We found that the eNpHR3.0-expressing mice did not spend significant time in the EtOH-paired compartment compared to eYFP expressing animals on Day 5 – in fact they appeared to show no preference whatsoever when the laser was activated (Figure 3e, ‘ON’). However, when animals were tested in the absence of laser activation of eNpHR, the level of CPP recall was fully intact (Figure 3e, ‘OFF’), suggesting that the EtOH preference memory had been formed during conditioning, but was unable to be expressed when the activity of Thy1+ cells is inhibited. Such observations, together with their strong EtOH-induced activation, suggested to us that BLA Thy1+ neurons are causally involved in both the associative *learning* and *recall* for alcohol rewards.

### BLA Thy1+ axonal activity differentially contributes to the acquisition and expression of alcohol CPP in a projection-specific manner

It has been well documented that BLA neurons project to many downstream regions, including the neuronal circuits for reward and fear ^23^. BLA projections to the NAcc and PFC have been implicated in reward-related behaviors, as distinct synaptic changes occur in neurons projecting to those targets with associative learning ^17, 24^. Our previous report also demonstrated that BLA Thy1+ neurons project to the canonical reward-related regions, including the NAcc and PFC, with minimal projections to CeA. To investigate if BLA Thy1+ neurons play a differential role in associative learning and memory in a projection-specific manner, we next targeted BLA Thy1+ axons in the NAcc vs. PFC (Figure 4a) and carried out optogenetic inhibition during Conditioning or Choice Testing, as we had previously done for the BLA Thy1+ cell bodies, above. We found that the inhibition of BLA-Thy1+ axonal activity in the NAcc and prefrontal cortex differentially prevented contextual place association and recall in CPP, respectively (Figure 4b-e). Specifically, we found that optogenetic inhibition of BLA Thy1+ axons in NAcc disrupted CPP formation (Figure 4c, ‘NAcc’), but did not significantly affect Choice-Test recall (Figure 4e, ‘NAcc’). In contrast, optogenetic inhibition (ON) of BLA Thy1+ axons in the PFC during Choice-Test disrupted CPP recall (Figure 4e, ‘PFC’), but did not significantly affect CPP formation (Figure 4c, ‘PFC’). These results suggest a double-dissociation in these pathways and functions, such that the BLA Thy1+ neurons differentially interact with downstream target areas during associative formation and retrieval of CPP memory.

**Figure 4.**
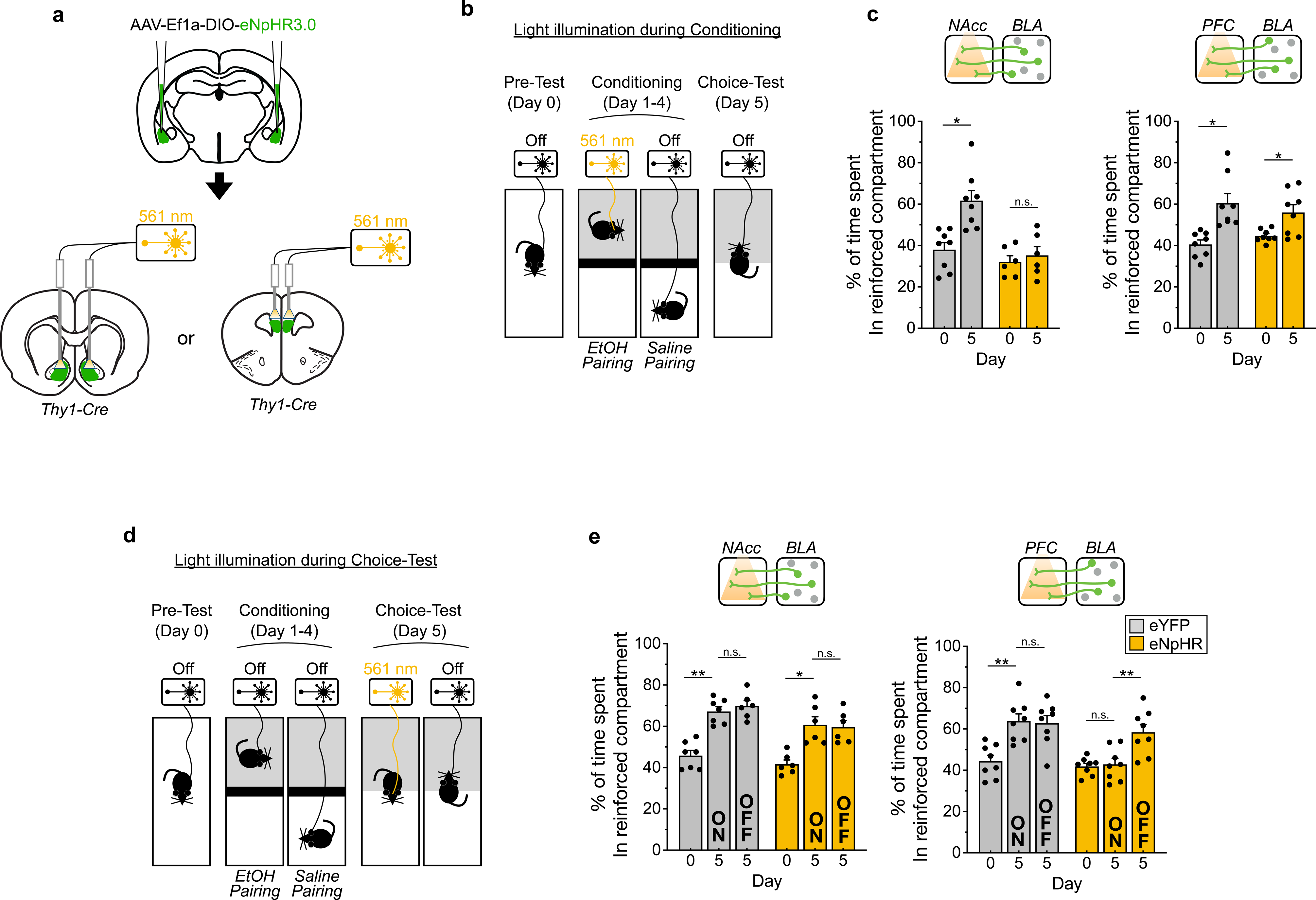
| BLA Thy1+ neuronal activity at axons in the NAcc and PFC during Conditioning and Choice-Test is necessary for CPP formation and recall, respectively. **a**, Scheme for Cre-dependent eNpHR/eYFP expression in BLA Thy1+ neurons and optogenetic manipulation at the BLA Thy1+ axons in NAcc or PFC. **b,** Illumination of light (561 nm, 15-20 mW) started 5 sec before EtOH injection and continued during the entire EtOH-pairing sessions on Conditioning Day 1-4. **c,** Optogenetic inhibition of BLA Thy1+ axons in the NAcc, not in the PFC, during Conditioning disrupted CPP formation. **d,** During Choice-Test, light illumination was carried out similarly for the entire session. **e,** Optogenetic inhibition (ON) of BLA Thy1+ axons in the PFC, not in the NAcc, during Choice-Test disrupted CPP recall. In a subsequent Choice-Test session without light illumination (OFF), mice injected with eNpHR displayed a similar level of CPP recall compared to mice injected with eYFP.

## Discussion

Addiction is often viewed as a disease of learning and memory ^2^. Here, we investigated whether alcohol could leverage neuronal circuits of natural learning and memory to lead to later alcohol-seeking behavior. We report that activity of Thy1-expressing excitatory BLA neurons correspond to repeated associative learning between alcohol administration and environmental context. We found an increased response of BLA Thy1+ neurons to alcohol over the course of conditioning, while the alcohol conditioning did not impact the activity of non-Thy1-expressing BLA neurons. The activity of this genetically defined neuron population also represented the alcohol-paired compartment by switching off their activity in the saline-paired compartment.

Next, we demonstrated greater activity from BLA Thy1+ neurons when entering a EtOH-paired compartment, whereas we saw decreased, or weaker, activity when exiting. Furthermore, we showed that inhibition of BLA Thy1+ neuronal activity at cell bodies during conditioning or choice-testing disrupted CPP formation or expression, respectively. Finally, we demonstrated that the acquisition and expression of alcohol CPP are differentially mediated by NAcc and PFC projections, respectively, from the BLA Thy1+ neurons. Together, this work reveals a sub-population of excitatory neurons in BLA differentially involved in acquisition and retrieval of alcohol-context association memory, raising the possibility of targeting specific subsets of amygdala neurons that encode memories of voluntary alcohol use.

Previous work has shown that reward-seeking behavior in response to conditioned reward-predicting cues relies on connectivity via glutamatergic changes within amygdala-striatal circuits. For example, BLA neuronal responses to cues has been shown to precede those of NAcc neurons, and cue-evoked excitation of NAcc neurons relies on BLA input ^25^. In addition, optical inactivation of direct projections from the BLA to the NAcc reduces behavioral responding for sucrose during the cue-reward pairing period in mice ^26^. Since the NAcc is involved in encoding of reward value and subsequent action selection that optimize reward seeking behaviors in response to sensory cues, the data show that the BLA, via excitatory and direct projections to the NAcc, contributes to motivated behavioral responding to obtain rewards. In addition to natural rewards, it has also been demonstrated that the exposure to alcohol, via prolonged heavy drinking, resulted in potentiation of glutamatergic activity in corticostriatal synapses ^27–29^. These changes in synaptic plasticity seem to sustain alcohol seeking and consequently promote habitual seeking behaviors. Similarly, an increase in extracellular levels of glutamate in the BLA and NAcc core has been observed during cue-induced reinstatement of alcohol seeking ^30^. Our current work provides a cell-type-specific substrate for some of these prior observed behaviors and physiological processes that deserves further examination.

The BLA is an integral hub not just for reward-related behavior but for regulating aversive learning and anxiety. It is involved in a number of behaviors relevant to aversion and stress, such as the acquisition, consolidation, and extinction of Pavlovian-conditioned fear memory ^15, 19^. The expression of fear memories and ultimately relapse depends on outputs of the subnuclei, including the BLA, of the amygdala ^31^. As the BLA is mainly composed of glutamatergic projection neurons, several reports demonstrated that a distinct subset of BLA neurons is involved in driving avoidance or approach behaviors based on their post-synaptic targets. These observations suggest that anatomically divergent and molecularly distinct populations of BLA neurons are involved in encoding positive or negative valence. Our results of the contribution of BLA Thy1+ neurons to alcohol CPP acquisition and recall indicate that these neurons are potentially involved in processing of positive valence. This is in addition to our prior work suggesting that this same population may serve as ‘Fear-off’ neurons that appear to suppress fear behavior and enhance fear extinction ^19, 20^. Thus, the same subpopulation of BLA neurons may both suppress fear behavior and support appetitive / positive valence-related behavior. Since valence is a subjective value and affected by the homeostatic needs of the animal, it would be interesting to investigate how prior experience of stress and repeated episode of heavy drinking affect this valence coding of BLA Thy1+ neurons in future studies.

Here we describe a genetically defined population of BLA neurons, Thy1+, that are activated by ethanol administration *in vivo*, and whose activation increases with repeated conditioning with ethanol. Optical inhibition of either the BLA Thy1+ neurons or their axons impairs alcohol conditioned place preference through differential projections to NAcc (necessary for conditioning) and PFC (necessary for recall). Importantly, as we previously demonstrated ^19^, the identification of cell-type-specific receptors or other molecular targets within BLA Thy1+ neurons may also lead to novel circuit-specific therapeutic targets for reducing conditioning or relapse of alcohol seeking behaviors in AUD.

In conclusion, this work highlights the BLA Thy1+ neuron population as a molecularly distinct, fundamental circuit in promoting the association of environmental cues and alcohol experience, and suggests that further examination of these neurons and their projections is relevant for the study of motivation to consume alcohol after abstinence.

## Methods

### Animals

All experiments were approved by and carried out in accordance with the Institutional Animal Care and Use Committee at McLean Hospital. All experimental and animal care procedures met guidelines outlined in the NIH Guide for the Care and Use of Laboratory Animals. C57BL/6J (B6) mice at 7 weeks of age were purchased from Jackson Laboratories (Bar Harbor, ME), and Thy1-YFP (Jackson Laboratory, Strain# 003782) and Thy1-Cre (Jackson Laboratory, Strain# 006143) were bred in B6 background and maintained in standard conditions under a 12:12-hour light/dark cycle (lights on: 07:00). Mice were group housed and allowed access to tap water and free (*ad libitum*) access to standard laboratory chow during the entire experimental period. After surgery, all the animals were single housed. For subsets of mice, blood samples (30-50 ul) were collected from the submandibular vein to measure blood ethanol concentration (BEC) using Analox Analyzer (Analox Instruments; courtesy of Klaus Miczek Laboratory).

### Conditioned Place Preference (CPP)

Mice were trained and tested in a CPP paradigm (Cunningham et al., 2006). A CPP box had 2 equal-sized compartments visually and tactilely different that were divided by an opaque guillotine door; the position of each compartment was counterbalanced. On day 0, mice were allowed to access both compartments for 20 min (Pre-Test), and the naturally preferred and non-preferred sides were noted. During Conditioning (day 1-4), mice were injected with 20% alcohol (v/v, 2 g/kg BW) and restricted to the non-preferred compartment or injected with saline and restricted to the preferred compartment for 15 min. A 2-trial-per-day (one for EtOH and the other for saline) procedure was used, where mice were randomly assigned to morning (10 AM) or afternoon (4 PM) group for the EtOH injection. During the Choice-Test on Day 5, mice were placed in the CPP apparatus without reinforcers (e.g., they were not given EtOH injections). Movement and locations of mice were collected with Ethovision software (Noldus).

### Immunohistochemistry

After behavioral testing, mice were deeply anesthetized with ketamine/xylazine (100 mg/kg, 10 mg/kg) mixture and transcardially perfused with PBS (pH 7.4), followed by paraformaldehyde (PFA, 4% in PBS). Brains were extracted, fixed overnight in 4% PFA, cryoprotected in sucrose (30% in PBS) for 48 hours, and sectioned into 50 um thick sections with a cryostat. Floating brain sections were treated in 3% H2O2 in PBS for 15 min, 50% EtOH in PBS for 30 min, blocked in 0.1% Triton-X/10% normal goat serum in PBS. The sections were incubated with Primary antibody against c-Fos (1:1000 rabbit anti-c-Fos, ProteinTech) overnight at 4 C, incubated with biotinylated secondary antibody (goat anti-rabbit, Vector laboratory, ABC kit (Vector laboratory) and TSA reagent (Perkin-Elmer). Finally, sections were counter-stained in DAPI, mounted on glass slides, and cover-slipped with mounting medium (Vectorshield, Vector laboratory). Images along the rostral-caudal axis of the BLA (AP -0.9 through -2.4 mm) were collected and c-Fos+ cells were quantified using CellProfiler.

### Viral constructs

AAV-Ef1a-DIO-GCaMP6f (1 x 10^13^ vg/ml) and AAV-Ef1a-DIO-eNpHR3.0 (1 x 10^13^ vg/ml) were purchased from the UPenn viral core and Addgene. All viruses were stored in aliquots at - 80C until use.

### Stereotaxic virus injection and optic fiber implantation

All surgery was performed under aseptic conditions and body temperature was maintained with a heating pad. Mice were anesthetized with a ketamine/xylazine (100 mg/kg, 10 mg/kg) mixture and placed in a stereotaxic apparatus. Ophthalmic ointment was applied to the eyes, hair was removed from the surface of the head with hair clippers, the area was scrubbed with alcohol and betadine, and an incision was made with a scalpel to expose the skull surface. After craniotomy, 250 nl of AAV-containing solution was injected into the BLA (ML ±3.2, AP -1.6, DV -5.3 mm). Then, for the optogenetic experiments, 2 optic fibers (200 um core, 0.48 NA, 5.0 mm length; RWD Life Sciences) were bilaterally placed and lowered to target the BLA. The scalp incision was closed with surgical sutures. For the calcium imaging studies, a second surgery was performed to implant an optic fiber (400 um core, 0.48 NA, 5.0 mm; RWD Life Sciences) to target the BLA after three weeks of recovery from the first surgery.

### Fiber photometry

Fiber photometry recordings were carried out between 4 and 8 weeks following viral vector injection. A fiber photometry system was used to measure bulk fluorescence from the BLA using a single optic fiber (1m, 400 um core, 0.48 NA; Doric Lenses) for both delivery of excitation light and collection of emitted fluorescence from GCaMP expressed in the BLA in real time. An RZ5P fiber photometry processor (Tucker-Davis Technologies, TDT) running Synapse software was used to drive two light-emitting diodes (LEDs) at 490nm ‘blue’ at 531 Hz, or at 405 nm ‘violet’ at 211 Hz (Doric lenses). These carrier frequencies were chosen to avoid cross-talk between channels. Excitation light from each LED was coupled into a 0.48NA, 400 um-core fiber, collimated, combined by a 425 nm long-pass dichroic mirror (minicube, Doric Lenses), and then was coupled into a 0.48 NA, low-autofluorescence 400 um patch cord (1 m, Doric Lenses). The far end of the patch cord was connected to implanted optic fibers with bronze sleeves (Doric Lenses). Emission light was collected via fluorescence photodetector heads integrated into the minicube. Signals were demodulated, digitized at 1 kHz, and recorded via the Synapse software. The start and end times of an EtOH CPP procedure were collected as transistor-transistor logic (TTL) time stamps using the digital input and output ports (mini-IO box, Noldus) and were then used to align the photometry signal and behavioral tracking data from EthoVision (Noldus). Both setups were tested for temporal precision to verify the synchrony of fiber photometry data and behavioral data. Photometry signals were analyzed using custom scripts as described in a previous report with modifications ^32^. Data from the signal and isosbestic control channels were extracted and smoothed using a local linear regression to reduce high-frequency noise from each signal. Next, a least-squares linear fit was applied to the 405 nm control signal to align to a 490 nm experimental signal to correct for photobleaching. Control and experimental signals were independently normalized to z scores, and the resulting z-scored reference signal was regressed onto and subtracted from the z-scored experimental signals to yield normalized experimental signals (zdFF). Fluorescent calcium events were detected in zdFF signals collected during conditioning sessions by calculating the zdFF for small chunks of data across the entire dataset with temporal windows of 1 second, centering and normalizing these windows, and then iterating this process for 100 times with temporally offset windows across the same data. The number of fluorescent calcium events that surpass a threshold (> 2.19) ^33^ during conditioning sessions were counted. Photometry signals during test sessions were aligned to transitions between compartments, and the window centered around each transition (−10 to 10 s) was extracted for analysis. Mean signals at transition (zdFF at time = 0 s) were analyzed. Subjects with lack of viral expression, incorrect fiber placement, or no change in preference for the EtOH-paired chamber were excluded from analyses.

### Optogenetics

Optogenetic excitation or inhibition of virally driven ChR2 and eNpHR3.0 was achieved using a 50 mW DPSS 473 nm laser (IkeCool Inc.) and a 150 mW DPSS 593 nm laser (RWD Inc.).

Patch cords were attached directly to chronically implanted fiber optics to the skull via a rotary joint (Doric Lenses) and suspended above a CPP apparatus. For the ChR2-mediated activation, animals received 10 mW light at 20 Hz (5 ms), whereas for the eNpHR3.0-mediated inhibition, animals received 20 mW tonic light.

### Quantification and statistical analyses

All statistical analyses were carried out using Prism (GraphPad) and in all cases, values showing p<0.05 were considered significant. All experimental data were subject to histological validation, where behavioral readouts were excluded if the conditions of the histological validation were not met (e.g. optical fibers were not correctly positioned above targeted brain areas).

## Acknowledgements

This work was supported by NIH awards R21-AA027450 (JS and KR), R01-AA030585 (JS), P50-MH115874 (KJR), R01-MH108665 (KJR), and the Frazier Institute at McLean Hospital (KJR).

